# Identifying neuron subtype-specific metabolic network changes in single cell transcriptomics of Alzheimer’s Disease using perturb-Met

**DOI:** 10.1101/2021.01.18.427154

**Authors:** Yuliang Wang

**Affiliations:** Institute for Stem Cell and Regenerative Medicine, University of Washington, Seattle, WA 98109, USA; Paul G. Allen School of Computer Science & Engineering, University of Washington, Seattle, WA 98195, USA

## Abstract

Metabolic aberrations are a prominent feature of Alzheimer’s Disease (AD). Different neuronal subtypes have selective vulnerability in AD. Despite the recent advance of single cell and single nucleus RNA-seq of AD brains, genome-scale metabolic network changes in neuronal subtypes have not been systematically studied with detail. To bridge this knowledge gap, I developed a computational method called perturb-Met that can uncover transcriptional dysregulation centered at hundreds of metabolites in a metabolic network. perturb-Met successfully recapitulated known glycolysis, cholesterol, and other metabolic defects in APOE4-neurons and microglia, many of which are missed by current methods. Applying perturb-Met on AD snRNA-seq data, I revealed that the four neuronal subtypes in the entorhinal cortex shows subtype-specific metabolic changes, namely mitochondrial complex I metabolism, ganglioside metabolism, galactose and heparan sulfate metabolism, as well as glucose and lipid metabolism, respectively. perturb-Met also revealed significant changes in protein glycosylation in the neuron subtype specifically found in AD brains. These subtype-specific metabolic changes may potentially underlie their selective vulnerability in AD. perturb-Met is a valuable tool to discover potential metabolic network changes in many other single cell or bulk transcriptomic studies.

## Introduction

Alzheimer’s Disease (AD) poses a significant disease burden, with 43.8 million people living with dementia globally, a situation that will further deteriorate with the world’s aging population[1]. Targeting metabolic dysfunction is a promising strategy to combat AD [2]. Genome-scale metabolic network models (GSMMs), which mechanistically connect thousands of metabolites and enzymes into a computable model, provide a global framework to systematically identify metabolic network changes. GSMMs have been successfully used to study many diseases, including AD[3,4]. However, existing applications of GSMM in AD are often based on bulk gene expression data of specific brain regions that contain multiple heterogeneous cell types[5] or rely on cell type-specific models manually curated from the literature[4], which may focus more on well-studied pathways. More importantly, AD does not affect all neurons to the same degree – some are more vulnerable than others, an observation known as selective neuronal vulnerability[6]. Earlier pioneering work showed that metabolic differences among neurons may provide a potential explanation[4]. However, the full extent of metabolic involvement in subtype-specific neuronal vulnerability in AD remains significantly under-explored and will be overlooked by GSMMs that use bulk expression data. Recently, massive amounts of single cell and single nucleus RNA-seq (sc/snRNA-seq) have been extensively used to measure transcriptomic changes in brain development, aging, and AD at the single cell level[7-10]. However, these datasets have not generated novel, high resolution insight into the neuron-subtype-specific metabolic dysfunctions in aging and AD.

To bridge this knowledge gap, we developed perturb-Met, which can identify transcriptional **perturb**ations centered at hundreds of **met**abolites in a genome-scale metabolic network (**Fig. 1**). perturb-Met adapts the Kullback-Leibler (KL) divergence concept from information theory to quantify relative changes in the distribution of transcript abundances for all genes connected to a metabolite between two conditions (e.g., AD vs. control). KL divergence quantifies how a statistical distribution differs from a second, reference distribution. perturb-Met is useful for both single cell and bulk transcriptomic studies.

**Figure 1.**
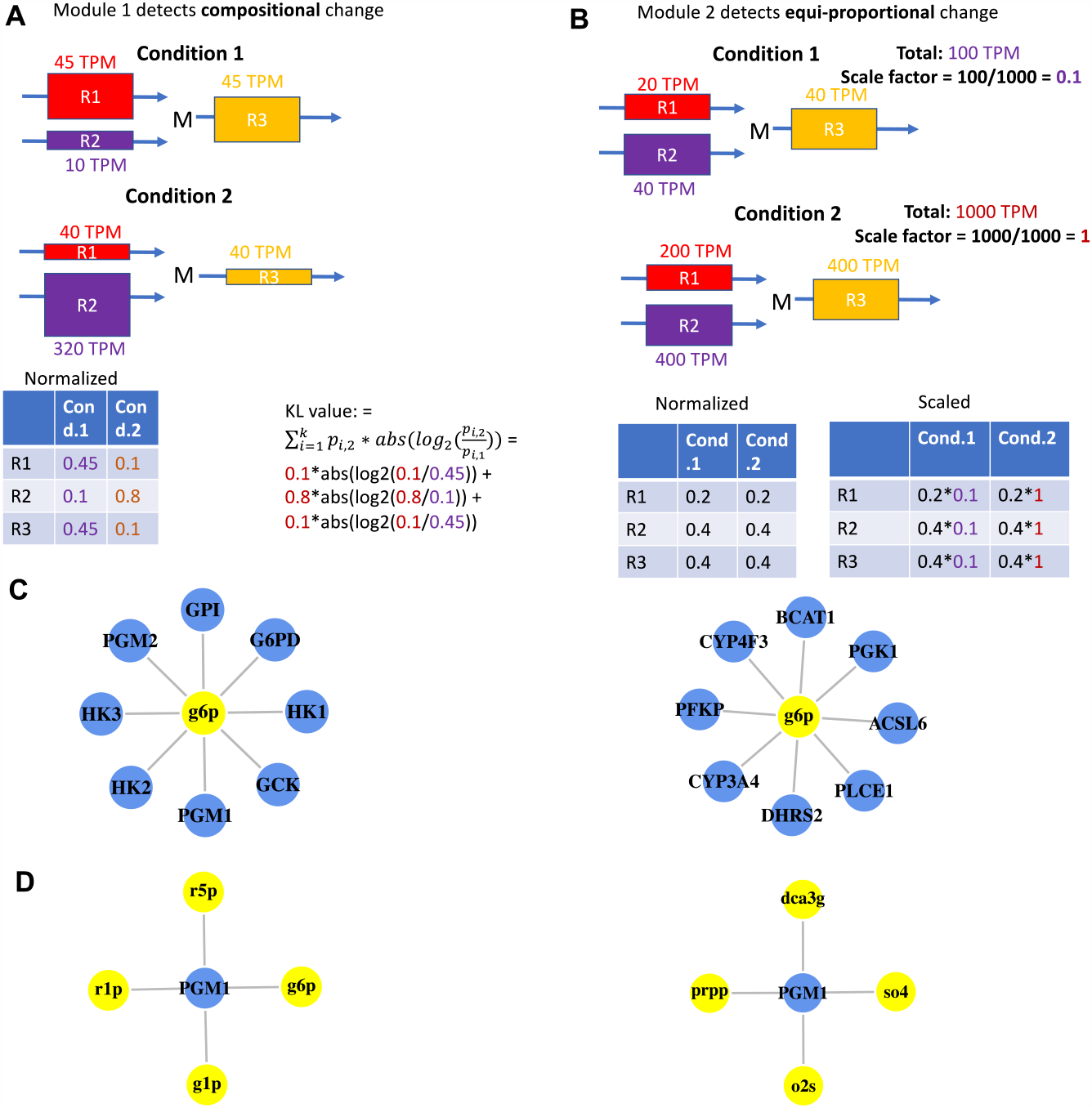
perturb-Met workflow. (**A)** Module 1 detects relative expression changes around a metabolite. Colored boxes with arrows represent metabolic reactions. Different colors denote different reactions. Size of the boxes is proportional to normalized enzyme mRNA levels that sum up to 1 for all genes connected to a metabolite. The metabolite M participates in R1, R2, and R3 reactions. R1 and R2 synthesize M, while R3 degrades M. Expression levels of the three reactions are shown in condition 1 (top) or condition 2 (bottom). A modified KL value formula is used to quantify relative expression changes. (**B**) Module 2 detects coordinated proportional expression changes around a metabolite. The condition with lower total expression level is scaled down before using the KL value formula. (**C**) Node-degree preserving permutation to calculate p-value. After permutation, each metabolite is still connected to the same number of genes, but the identities of those genes changed. (**D**) Similarly, each metabolic gene is still connected to the same number of metabolites, but the identities of those metabolites are randomly chosen.

perturb-Met offers several advantages over existing methods. First, the commonly used pathway-level enrichment analysis groups metabolic genes into distinct pathways, which does not account for the fact that metabolites often span and connect multiple pathways. Moreover, pathway-level analysis often lacks sufficient resolution: transcriptional changes may be concentrated in only a few metabolite and involve a small number of key enzymes within a pathway. perturb-Met addresses both problems by considering relative changes of transcript abundances at individual metabolites. Further, although Reporter Metabolite analysis can also identify transcriptional dysregulation around metabolites instead of pathways[11], it still relies on univariate differential expression analysis for each gene separately. This suffers from two potential problems. First, because it asks the question, *Is there more extensive differential expression than expected by chance?*, it often requires a minimum and a maximum number of genes connected to a metabolite (e.g., minimum 3 genes, maximum 50 genes[12]), which may miss important metabolites connected by only two genes (examples in **Fig. 2D** and **E**). Second, even if more than 3 genes are connected to a metabolite, Reporter Metabolite analysis would still require enough of them to be significantly differentially expressed. A metabolite may not be selected if only one of its connected genes is differentially expressed. However, using perturb-Met, it is still considered noteworthy if the magnitude of expression change for one gene significantly alters the relative distribution of transcript abundances of all genes around a metabolite (example in **Fig. 3F**).

**Figure 2.**
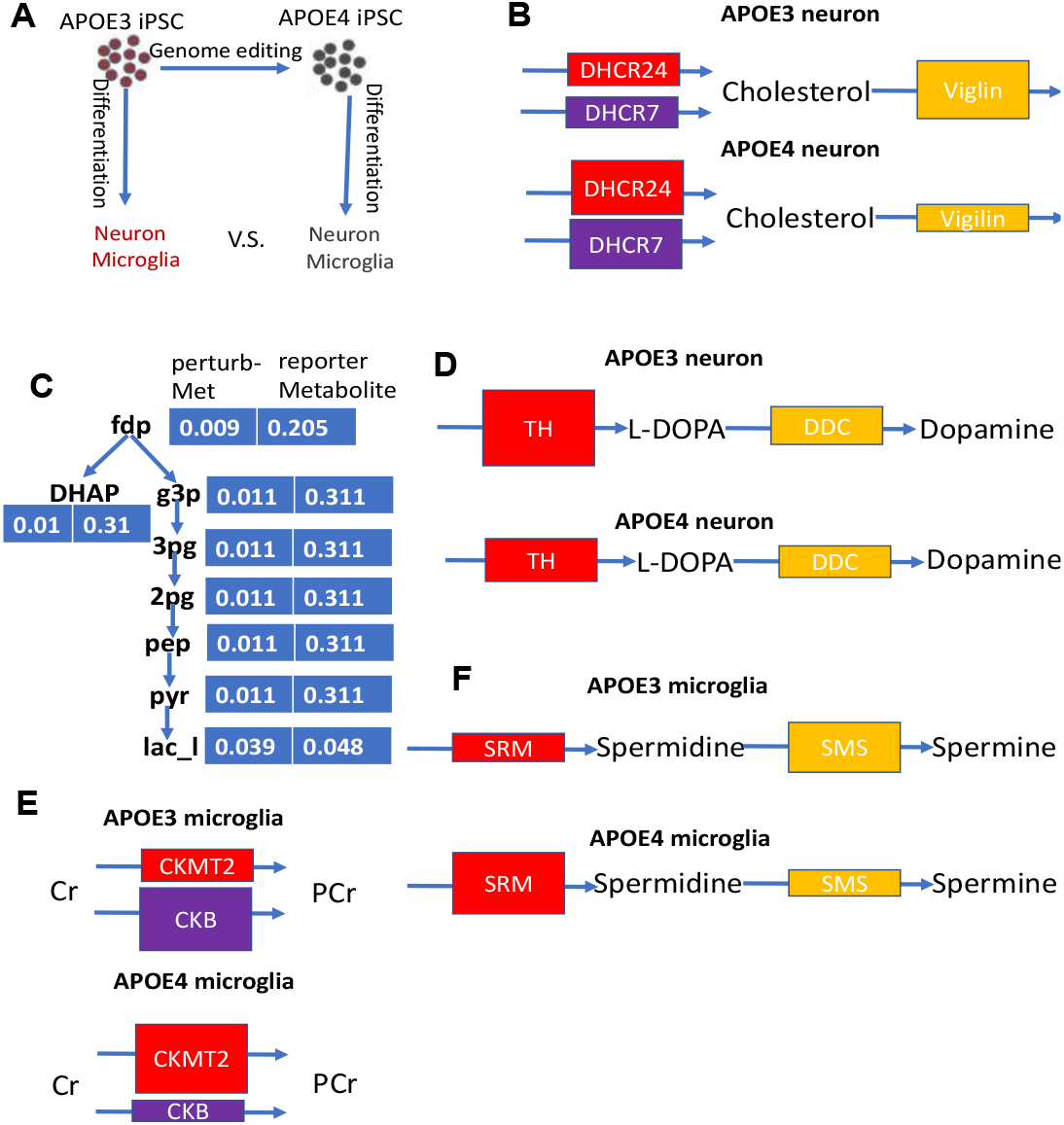
Validation of perturb-Met on APOE-4 vs. APOE-3 neurons and microglia. (**A)** CRISPR genome editing was used to create isogenic iPSCs that carry either APOE3 (control) or APOE4 (pathogenic) variants. These iPSCs are differentiated into neurons and microglia, which are then compared in this study. (**B**) perturb-Met identified cholesterol metabolism as being different between APOE4 and APOE3 neurons due to changes in the relative expression levels of cholesterol synthesis and transport enzymes. Colored boxes with arrows represent metabolic reactions. Different colors denote different reactions. Size of the boxes is proportional to enzyme mRNA levels. (**C**) perturb-Met recapitulated metabolic changes in glycolysis in APOE4 neurons. **Bold text** indicates metabolites in glycolysis; blue boxes adjacent to each metabolite are p-values from perturb-Met and Reporter Metabolite analysis, respectively. Blue arrows represent individual steps in glycolysis. (**D**) L-DOPA metabolism in APOE4 neurons is identified by perturb-Met as potentially interesting due to relative expression changes of tyrosine hydroxylase (TH) and dopa decarboxylase (DDC). (**E**) Creatine/phosphocreatine metabolism may be perturbed in APOE4 microglia due to relative expression changes of mitochondrial (CKMT2) and cytosolic (CKB) isoforms of creatine kinase. (**F**) Spermine/spermidine metabolism identified by perturb-Met in APOE4 microglia due to relative expression changes of spermidine synthase (SRM) and spermine synthase (SMS). fdp: Fructose 1,6-Bisphosphate; DHAP: Dihydroxyacetone Phosphate; g3p: Glyceraldehyde 3-Phosphate; 3pg: 3-Phospho-D-Glycerate; 2pg: 2-Phospho-D-Glycerate; pep: Phosphoenolpyruvate; pyr: Pyruvate; lac-l: L-lactate.

**Figure 3.**
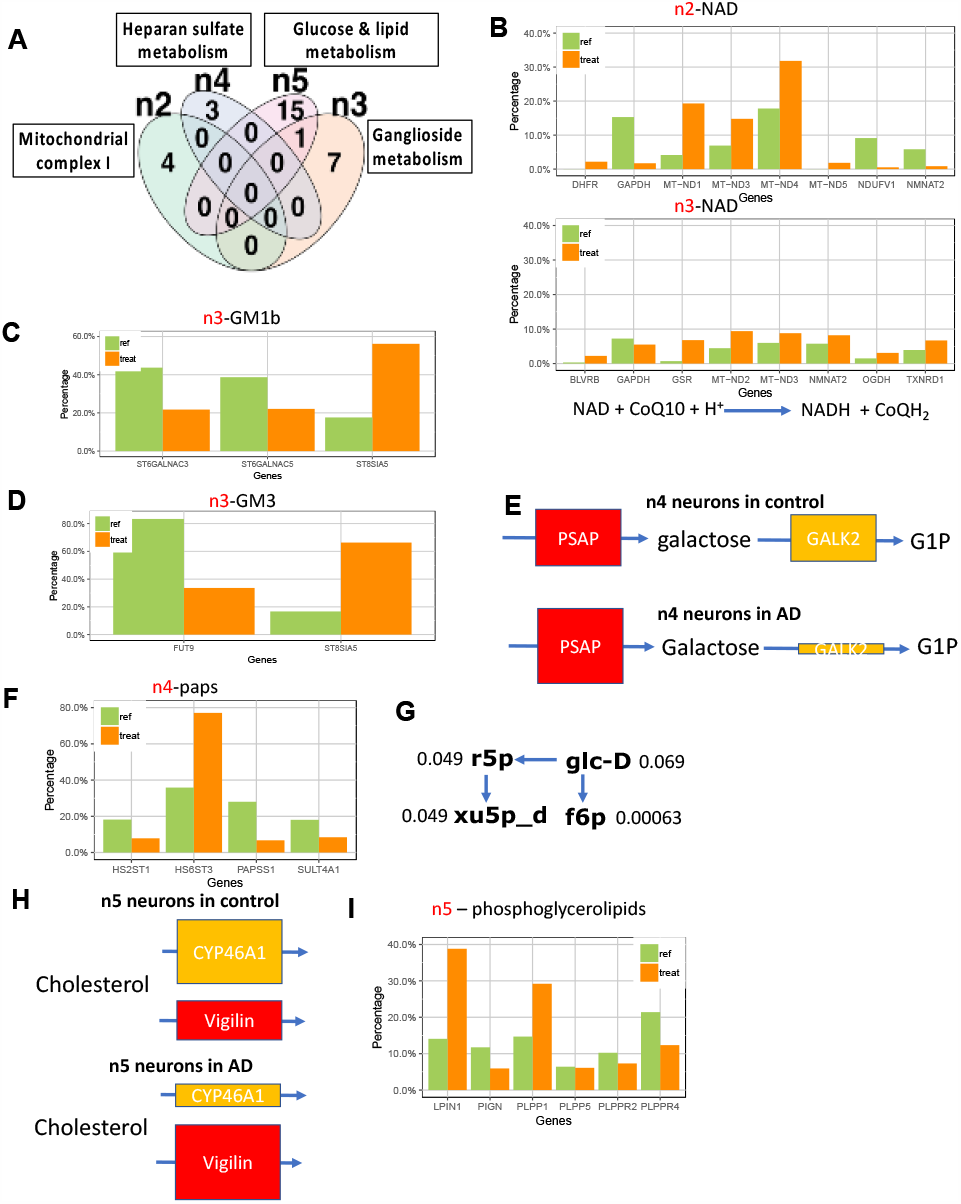
perturb-Met identified neuron sub-cluster-specific metabolic changes in response to Alzheimer’s disease (AD). **(A)** Venn diagram of metabolites identified by perturb-Met in each neuron sub-cluster showed that each sub-cluster had unique metabolic responses. **(B)** Bar plot of metabolic genes connected to NAD in n2 sub-cluster (top) and n3 sub-cluster (n3) showed NAD metabolism uniquely changes in n2. Expression values are normalized such that all enzymes connected to a metabolite sum to 100%. Green bars: normalized expression values in control neurons; red bars: normalized expression values in AD neurons. For clarity, only metabolic enzymes showing noticeable changes are plotted. **(C, D)** Enzymes in gangliosides GM1b (C) and GM3 (D) metabolism show relative expression changes in n3 neurons. **(E)** Galactose metabolism is perturbed in the n4 sub-cluster due to relatively decreased galactose kinase (GALK2) expression and increased Prosaposin (PSAP) expression. **(F)** There is a potentially increased transfer of sulfate from paps (3’-Phosphoadenosine-5’-phosphosulfate) to heparan sulfate due to increased expression of HS6ST3 in n4 AD neurons. **(G)** 4 metabolites in glucose metabolism are identified as perturbed in n5 neurons in response to AD. **(H)** Cholesterol is identified as perturbed in n5 neurons due to relative expression changes of CYP46A1 and vigilin. **(I)** Multiple phosphoglycerolipids are identified as perturbed in n5 neurons due to increased expression of lipin 1 (LPIN1) and phospholipid phosphatase 1 (PLPP1) genes in AD.

We validated the performance of perturb-Met by showing that it successfully recapitulated known cholesterol, glycolytic, and other metabolic dysfunctions in neurons and microglia harboring the e4 allele of APOE, the strongest genetic risk factor for late-onset AD [13], which are missed by existing methods. We then used perturb-Met to identify neuron subtype-specific metabolic changes in AD using snRNA-seq[10], recapitulating many literature-supported metabolic changes and uncovering new ones for further experimental validation. Specifically, we found that the four neuronal subtypes identified in a recent scRNA-seq of human AD[10] are each characterized by unique metabolic dysfunctions in mitochondrial complex I metabolism, ganglioside metabolism, galactose and heparan sulfate metabolism, as well as glucose and lipid metabolism. Comparing the AD- and control-specific neuron subtypes revealed changes in protein glycosylation, known to be important for many key AD genes [14]. Our results provide potential metabolic explanations for selective neuronal vulnerability in AD and may offer more precise guidance to target metabolic dysfunctions in this disease. Our perturb-Met method can also be applied to uncover potential metabolic reprogramming in a wide range of human diseases and developmental processes based on either single cell or bulk transcriptomics data.

## Results

### 1 Overview of the perturb-Met method

perturb-Met combines results from two computational modules. The first module detects relative compositional changes in transcript abundances across genes (**Fig. 1A**): gene B becomes more dominant in condition 2 (e.g., 10% of total transcript abundance in condition 1 to 80% of total abundance in condition 2). The second module detects cases where the relative expression levels do not change, but total expression levels change (**Fig. 1B**). In each module, we calculate the adapted KL divergence and use 1000 permutations to derive a p-value (Methods). Permutation p-value is defined as the fraction of times that the expression changes of randomly sampled sets of genes (where each metabolite is still connected to the same number of genes and each metabolic gene is connected to the same number of metabolites, **Fig. 1C & D**) have the same or larger KL divergence than the actual metabolic gene set based on metabolic network topology. We then use Fisher’s method to combine the p-values from modules 1 and 2 to obtain a single p-value for each metabolite[15]. This p-value is then used to rank metabolites.

In contrast, Reporter Metabolite analysis first calculates a p-value for each gene separately and then uses inverse normal distribution to combine the p-values for all genes around a metabolite into a single score; it also uses permutation to derive a p-value for the combined score[11]. perturb-Met differs from Reporter Metabolite analysis in that it considers expression changes in each metabolic gene together with the expression levels of all other metabolic genes consuming/producing the same metabolite, instead of testing differential expression for each gene separately. For sc/snRNA-seq data, perturb-Met also performs cell label permutation, where the cluster (1, 2, 3, etc.) and phenotype (e.g., AD, control) are randomly assigned (Methods).

### 2 Validation of perturb-Met in APOE4-neurons and microglia

To evaluate the performance of perturb-Met and compare it with Reporter Metabolite analysis [11], we applied both methods on a dataset with well-characterized underlying metabolic changes.

The APOE4 variant is a major genetic risk factor for AD[16] and is known to cause aberrations in cholesterol and glucose metabolism in neurons[13]. We applied both perturb-Met and Reporter Metabolite analysis on a bulk RNA-seq dataset of APOE4-(pathogenic) and isogenic APOE3-(control) neurons[17] (**Fig. 2A**). perturb-Met correctly identified cholesterol as perturbed between APOE4-vs. APOE3-neurons (**Fig. 2B**). perturb-Met also identified significant transcriptional dysregulation around many glycolytic metabolites (**Fig. 2C**).

Both cholesterol and glycolytic metabolites are missed by the Reporter Metabolite analysis: this method requires a large number of the genes connected to a metabolite to be significantly differentially expressed in order to detect a metabolite as perturbed, and it requires a metabolite to be connected to at least three genes in order to perform significance testing. perturb-Met does not have such restrictions.

In addition to well-known aberrations in cholesterol and glucose metabolism, we identified other literature-supported changes in APOE4-neurons that are missed by Reporter Metabolite analysis (**Table 1**, full list in Table S1). Interestingly, we identified transcriptional dysregulation in the synthesis of neurotransmitters dopamine and serotonin. perturb-Met found tyrosine and L-DOPA to be perturbed mainly due to decreased expression of tyrosine hydroxylase in APOE4-neurons, which converts tyrosine into L-DOPA (**Fig. 2D**); L-DOPA is a precursor for dopamine. Reduced dopamine transporter and tyrosine hydroxylase levels have been previously observed in the nucleus accumbens of AD patients[18], and augmenting dopamine transmission has been shown to improve cortical neurotransmission and cognitive performance of AD patients[19]. Similarly, perturb-Met also found several metabolites in serotonin synthesis (tryptophan, 5-hydroxy-tryptophan and serotonin) due to decreased DDC (DOPA decarboxylase) expression. This could lead to reduced serotonin production by APOE4 neurons compared to APOE3 neurons. Increasing serotonin signaling has been shown to reduce amyloid-β in mouse models of AD and in human[20] and to improve cognitive performance of AD patients[21]. Our analysis therefore suggests that adding serotonin and/or dopamine in APOE4-neurons may confer beneficial effects.

**Table 1.**
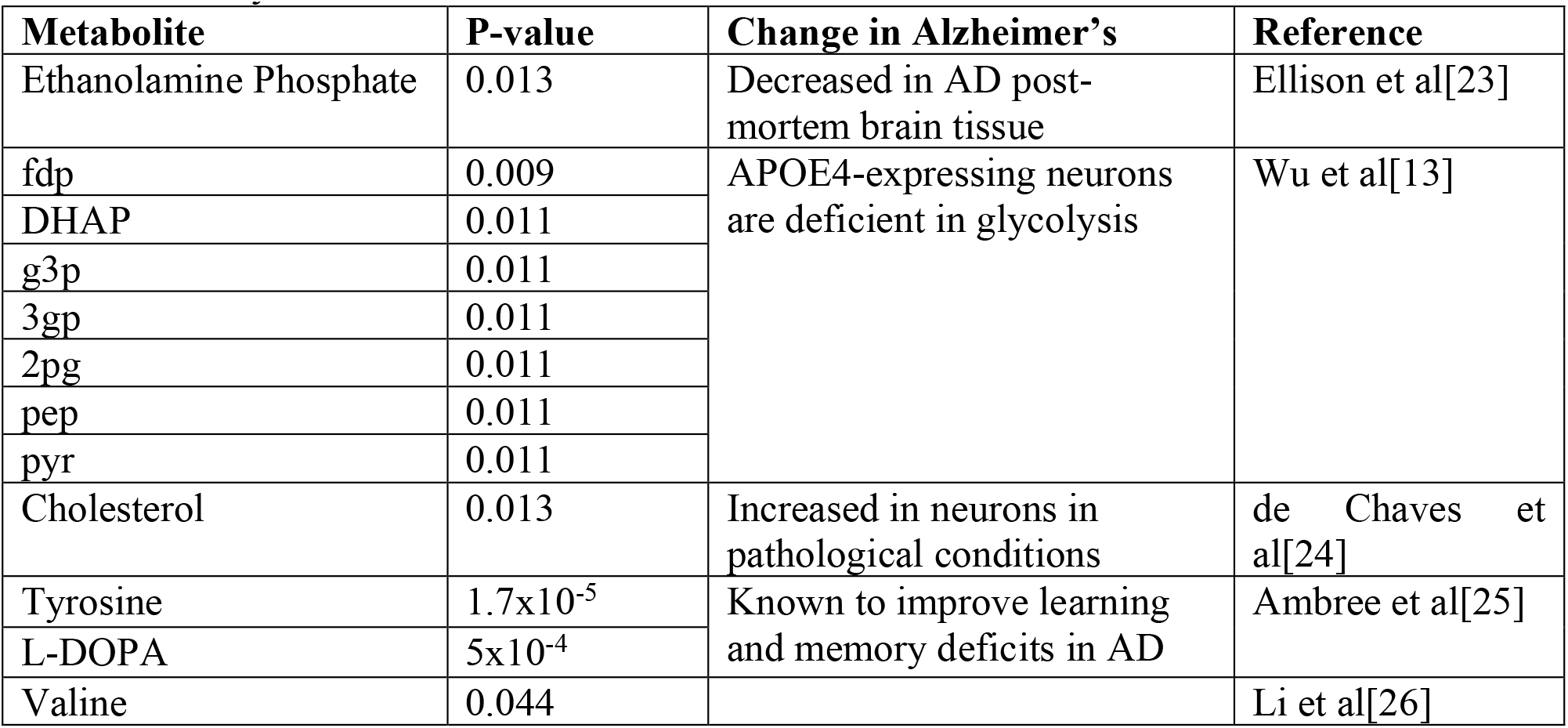
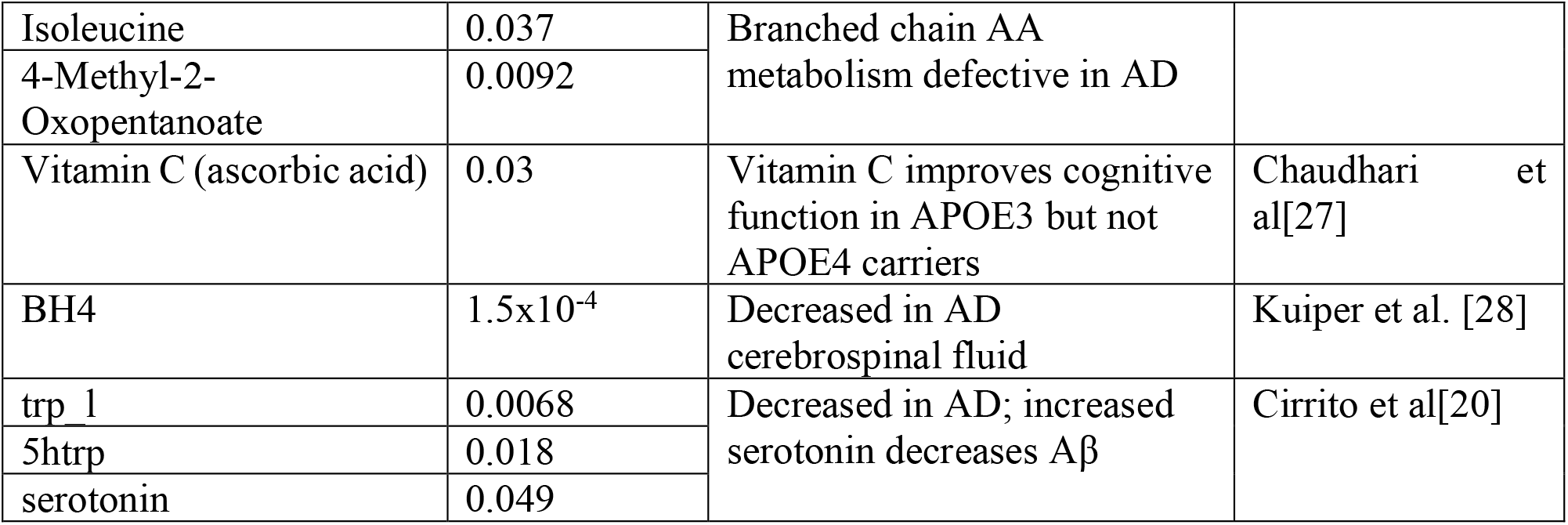
Literature-supported metabolites identified by perturb-Met but missed by Reporter Metabolite analysis in the APOE4-neuron.

Besides metabolites with existing literature support listed in **Table 1**, we also identified many novel metabolites for future experimental validation. One example is multiple components of the glycine cleavage system (alpam, alpro, lpro, dhlpro): increased glycine accumulation can cause motor dysfunction[22], but its role in AD is unclear. Additionally, both Reporter Metabolite analysis and perturb-Met identified multiple metabolites in the one-carbon metabolism as perturbed (mlthf, dhf, thf, Table S1). This suggests that defective one-carbon metabolism may be an important phenotype in APOE4-associated neuronal metabolic dysfunction.

Besides neurons, metabolic aberrations in microglia are increasingly recognized as being important for AD pathogenesis[29]. Therefore, we focused on the effects of the APOE4 variant on microglia, as well. Table 2 lists metabolites identified by perturb-Met but missed by Reporter Metabolite analysis (full list in Table S2).

**Table 2.** Literature-supported metabolites identified by perturb-Met but missed by Reporter Metabolite analysis in APOE4-microglia.

| Metabolites | P-value | Change in AD | Reference |
| --- | --- | --- | --- |
| Phosphocreatine | 0.0029 | Significant increase in entorhinal cortex of APOE4 aged mice | Area-Gomez et al[30] |
| S-Adenosylmethionine | 0.01 | Known to induce microglia activation and neuronal damage | Seminotti et al[33] |
| Spermine | 0.0071 | Has anti-inflammatory effects on microglia | Choi & Park[34] |
| Spermidine | 0.0076 |  |  |
| Uroporphyrinogen III | $6.3 \times 10^{-4}$ | Induces reactive microglia | Lin et al[35] |
| Protoporphyrin | 0.0073 |  |  |
| Protoheme | 0.038 |  |  |
| Methylmalonyl Coenzyme A | 0.001 | Known to induce microglia activation | Gabbi et al[36] |
| Succinate | 0.04 | Known to be pro-inflammatory | Mills et al[37] |

Phosphocreatine is a rapidly mobilizable reserve of high-energy phosphates that has previously been reported to be highly increased in the entorhinal cortex of APOE4 mice[30]. Our analysis further infers that microglia may contribute to increased phosphocreatine. Phosphocreatine is prioritized by perturb-Met due to a dramatic shift of the relative expression of cytosolic (CKB) vs. mitochondria creatine kinase (CKMT2) (**Fig. 2E**). Interestingly, the APOE4 variant is known to increase production of nitric oxide in microglia [31], which has been shown to inactivate CMKT2 (Sarcomeric Mitochondrial Creatine Kinase) and cause compensatory up-regulation of its expression, both in absolute levels and relative to the cytoplasmic CKB[32]; these literature results support the scenario in **Fig. 2E**.

perturb-Met also recovered several metabolites known to induce or repress microglia activation. S-adenosylmethionine (SAM), heme, methylmalonyl CoA and succinate are all known to promote inflammatory activation of microglia or macrophages by increasing reactive oxygen species, while spermidine (**Fig. 2F**) has anti-inflammatory effects on microglia by blocking NF-κB, PI3K/Akt and MAPKs signaling pathways (**Table 2**). This suggests that an APOE4 variant may influence microglia immune phenotypes by perturbing their metabolic homeostasis.

We also found novel metabolites not previously linked to microglia. The top prediction is N-Acetyl-D-Mannosamine (p-value 2.82×10^−6^), shown to improve cognitive function in aged mice[38], but its specific function in microglia is unknown. We also found multiple phospholipid species (Table S2). Phospholipids are known to influence microglia morphology[39]. Further, we found 11-cis-retinol (p-value 0.014) and 11-cis-retinal (p-value 0.019). Although 9-cis retinoic acid has been shown to suppress inflammatory response in microglia[40], the role of 11-cis retinol has not been studied and may be worth investigating. None of these metabolites are identified by Reporter Metabolite analysis.

In summary, compared to Reporter Metabolite analysis, perturb-Met uniquely identified many known, literature-supported metabolites as transcriptionally perturbed between APOE4-vs. APOE3-neurons and microglia. The novel metabolites predicted by perturb-Met provide new potential targets to correct metabolic dysfunctions in APOE4-neurons and microglia.

### Neuron subtype-specific analysis reveals unique metabolic expression changes in AD

Existing metabolic network models of AD are based on the transcriptomes of specific brain regions[5], which may miss extensive heterogeneity within the same cell type or across different cell types within the same region. This is important, because certain types of neurons are more susceptible to AD [6], and genome-scale metabolic networks have already been used to identify potential metabolic differences between neuron subtypes that may explain selective neuronal vulnerability in AD[4]. However, due to the smaller scale of these models, the extent of cell subtype-specific metabolic differences remains under-explored. We therefore used perturb-Met to identify metabolic heterogeneity for multiple subtypes of neurons.

I followed the subtype identified in Grubman et al. [10] and performed cell sub-cluster-specific perturb-Met analysis for neurons. We focused on neuron sub-clusters n2, n3, n4 and n5, which contained significant numbers of neurons from both AD and control brain samples. We also filtered out metabolic genes expressed in less than 10 cells in both AD and control samples. In addition to network permutation (**Fig. 1 C** and **D**), we also performed cell label permutation to ensure that observed relative expression changes were not due to random noise in the expression data (Methods). I found that neuron subtypes show largely unique metabolic perturbations in AD, with only a few shared between two neuron subtypes (**Fig. 3A**). Full results are in supplemental tables S3-S6.

Neuron sub-cluster n2 consists of excitatory neurons from cortical layers II to VI and is selectively vulnerable in AD. perturb-Met identified NAD/NADH and co-enzyme Q10 due to higher mitochondrial complex I expression (MT-ND1, ND3, and ND4, **Fig. 3B**). Interestingly, other complexes of the mitochondrial OXPHOS machinery did not show increased expression in AD.

This complex I up-regulation is also unique to n2 since other sub-clusters such as n3 neurons showed little change. Mild inhibition of complex I activity is known to restore energy homeostasis and avert cognitive decline in animal models of AD [41,42].

Neuron sub-cluster n3 is a mixture of NDNF+ and CCK+ interneuron subsets [10]. CCK+ interneurons shown Aβ accumulation and its number declines with age in AD [43]. 4 out of the 8 metabolites identified by perturb-Met are involved ganglioside metabolism: GM3, GD1a, GM1b, and CMP-sialic acid (**Fig. 3C** and **D**). The relevance of our prediction is supported by literature. Gangliosides concentration decreased in AD [44]. Recent studies demonstrate that physiological concentrations of ganglioside GM1 inhibits Aβ oligomerization [45], and ganglioside GQ1b reduces Aβ plaque deposition and tau phosphorylation and ameliorates cognitive deficits in the triple-transgenic AD mouse model (3xTg-AD) [46]

Neuron sub-cluster n4 corresponds to PVALB+ and SST+ layer IV–VI interneurons. This subtype of interneurons are impaired in AD [47,48]. The top perturbed metabolite is galactose due to decreased expression of galactokinase 2 (**Fig. 3E**). Interestingly, chronic exposure to galactose has been shown to induce neurodegeneration and increase oxidative damage [49]. We also identified altered heparan sulfate metabolism (paps), mainly driven by increased expression of HS6ST3 (**Fig. 3F**). Heparan sulfate proteoglycans are required for cellular uptake of tau [50]. Further, we identified uracil due to UPP2, which catalyzes the interconversion between uracil and uridine. Uridine is known to protect neurons from glucose-deprivation-induced cell death, and UPP2 is key for this protective effect [51].

The n5 sub-cluster includes layer I, II, and VI VIP+ interneurons, which have not been shown to be impaired in AD [52]. We found multiple glycolysis metabolites as different between AD and control conditions (**Fig. 3G**), for example, f6p (fructose-6-phosphate) was identified due to increase in hexokinase 1 in AD. Besides changes in glucose metabolism, we also detected changes in cholesterol metabolism due to increased HDLBP (aka. vigilin) expression (**Fig. 3G**). Vigilin is known to regulate lipid metabolism[53] and co-localizes with APOE [54], and its expression is actually lower in APOE4 neurons (**Fig. 2B**). Further, we observed multiple phosphoglycerolipids due to increased LIPIN1 and PLPP1 expression in AD (**Fig. 3H**), which are involved in phosphatidate (PA) and diacylglycerol (DG) conversion. Lipin 1 expression is up-regulated by injury, and its inhibition is known to promote axon regeneration [55].

In summary, perturb-Met revealed neuron subtype-specific changes, many of which are known to be important in AD. The metabolites and enzymes uniquely perturbed in each neuron subtype may provide important clues on selective neuronal vulnerability in AD.

### AD-specific neuron sub-cluster shows dysregulated protein glycosylation

In addition to n2-n5 neuron sub-clusters, which contain significant numbers of neurons from both AD and control samples (27% − 58% neurons from AD), there are two additional clusters: n1 and n6, where 91% and 98% of all neurons come from AD and control samples, respectively[10]. We found multiple metabolites in glycan metabolism to be different between AD- and control-specific neuron sub-clusters---UDP, UDP-glucuronate, UDP-galactose, UDP-GlcNAc, and heparan sulfate---due to expression changes in multiple enzymes, such as CHSY3, B3GAT1, B4GALNT4, FUT9, MGAT4C, HS6ST3, etc (**Fig. 4**, full results in Table S7). The aberrant protein glycosylation is an important feature of AD [56]. In particular, UDP-GlcNAc (uridine diphosphate N-acetylglucosamine) is an important substrate for protein glycosylation. Interestingly, we found that in AD-specific n1 neurons, there is increased β1,4 -N-glycosylation due to increased MGAT4C and decreased β1,6 -N-glycosylation (**Fig. 4A** and **B**). Many key AD-related genes are N-glycosylated, including APP, BACE1, and TREM2 [14]. In particular, N-glycan structures of APP influence its transport and secretion [14]. However, the effects of specific N-glycan structures on AD-related proteins remain unclear. We also observed (1) an increased transfer of UDP-GlcNAc and UDP-glucuronate to chondroitin sulfate due to increased CHSY3 and B3GAT1, and (2) a decreased incorporation into heparan sulfate due to lower EXT1 expression in AD (**Fig. 4C** and **D**). Chondroitin sulfate proteoglycans, found near senile plaques and neurofibrillary tangles [57], are known to inhibit neurite outgrowth after injury, and blocking chondroitin sulfate restores memory in tauopathy-induced neurodegeneration [58]. We also identified relative shifts of O-vs. N-sulfation of heparan sulfate in AD vs. control-specific neurons (**Fig. 4 E** and **F**). Heparan sulfate is also identified in cluster n4 AD vs. control comparison, and heparan sulfate proteoglycans are required for cellular uptake of tau [50].

**Figure 4.**
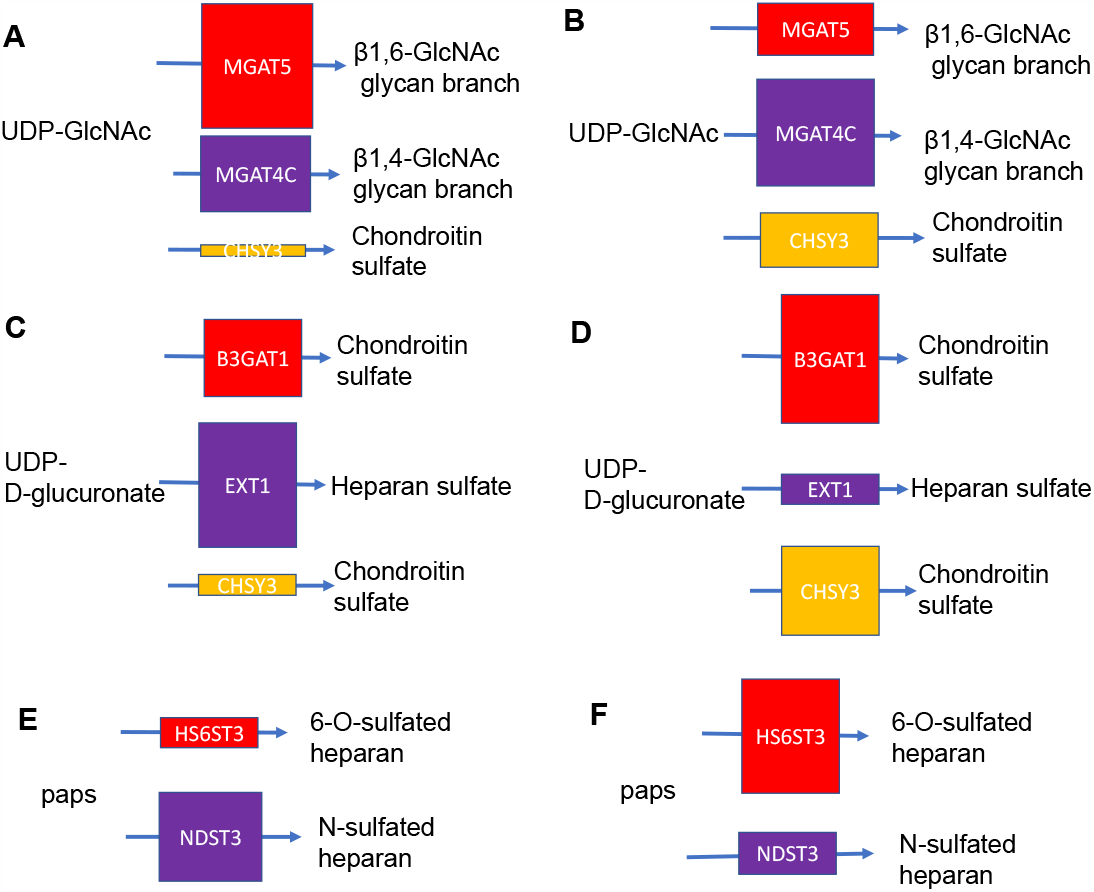
Glycan metabolism differs between the AD-specific n1 neuron sub-cluster and the control-specific n6 neuron sub-cluster. **(A**,**B)** Uridine diphosphate N-acetylglucosamine is a key substrate for multiple glycan modification reactions, and its transfer into different types/structures of glycans differs between control (A) and AD (B). Similarly, UDP-glucuronate transfer into different glycan species differs between control **(C)** and **(D)**, with increased transfer into chondroitin sulfate and reduced transfer into heparan sulfate in AD. **(E**,**F)** There is increased paps (3’-Phosphoadenosine-5’-phosphosulfate) incorporation into heparan sulfate via 6-O-sulfation than N-sulfation in AD (F) compared to control (E).

### Neuron subtype-level analysis revealed potential false positive and false negative discoveries due to bulk averaging

Comparing FACS-sorted neurons in AD vs. control conditions is a typical way to identify cell type-specific changes in AD [59]; however, expression profiles from these studies represent the average expression profiles of all purified neurons. Therefore, we tested whether the metabolites identified by sub-cluster-specific analysis in Fig. 4 are also captured by comparing the average expression profiles of all neurons in AD vs. the control. Among the 32 significant metabolites (p-value <0.05) perturb-Met identified on the average profile of all neurons (Table S8), 8 (25%) were also found in one of the four subtypes. The other 24 were found to be significant by bulk averaging but not in any one of the neuronal subtypes. On the other hand, there are 22 metabolites identified as significant by perturb-Met in at least one neuron subtype but not by bulk averaging (**Fig. 5A**).

**Figure 5.**
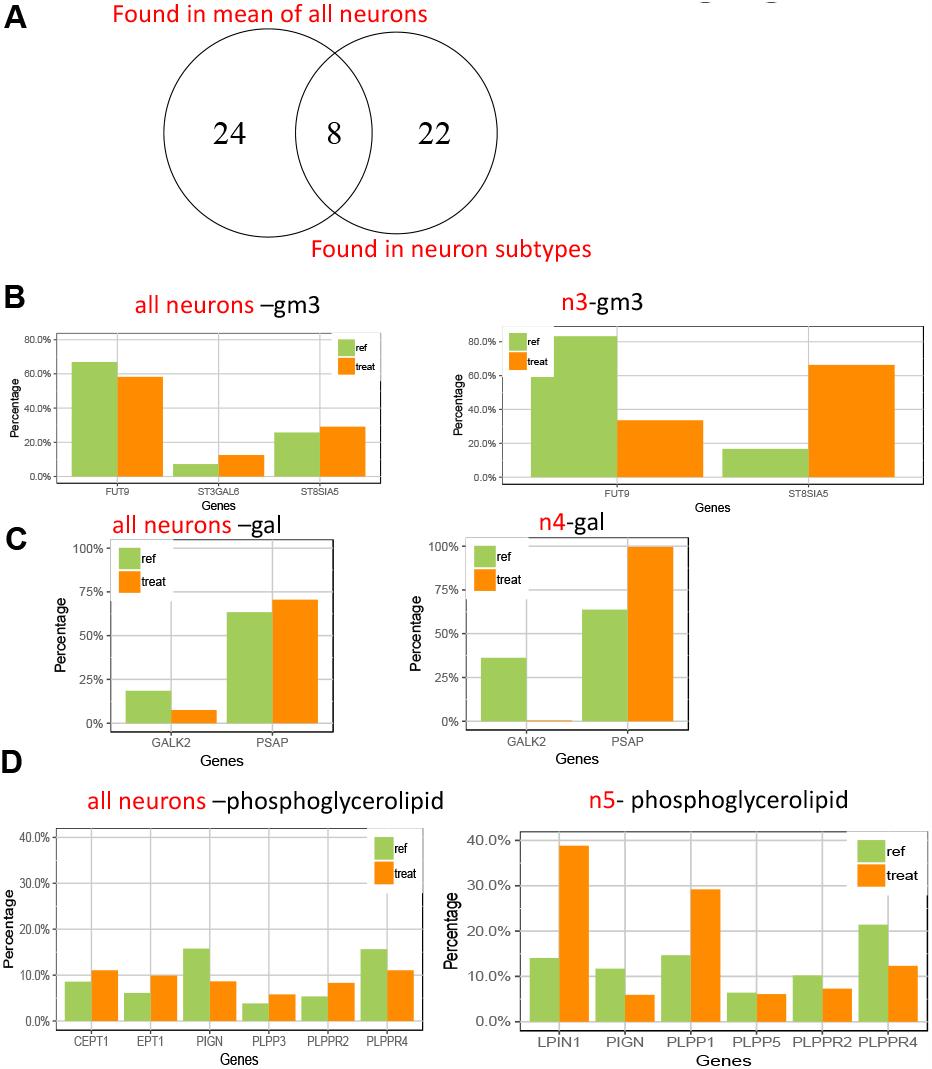
A. Applying perturb-Met on the average expression of all n2-n5 neurons in AD vs. the control, or applying it on each sub-cluster separately, resulted in different sets of metabolites identified as perturbed. (B, C, D) Three metabolites that are identified as perturbed in n3, n4 and n5 sub-clusters but not when using the average expression profiles of all neurons. Left: relative expression pattern in average expression profiles of all neurons in AD or control; right: relative expression pattern in specific neuron sub-clusters. Not all enzymes are shared because some are not plotted due to their low expression or small expression changes.

Some interesting metabolites detected in each neuron sub-cluster---GM3 in n3 (**Fig. 5B**), galactose in n4 (**Fig. 5C**), and glycerophospholipid in n5 (**Fig. 5D**)---are not detected using bulk averaging of all neurons in AD vs. the control. This is likely because expression changes occur only in a neuron sub-cluster, and bulk averaging is not sufficiently sensitive to pick up these changes.

There are also many metabolites are identified by bulk averaging but not by sub-cluster-specific analysis: genes connected to these metabolites are expressed in more than 10 neurons across all sub-clusters but in less than 10 cells in any specific sub-cluster in both AD and control conditions; thus, they are not considered in sub-cluster-specific analysis. We hypothesize that since these genes are not expressed in substantial numbers of cells in any sub-cluster, their expression levels are likely noisy and may not be biologically interesting. This result demonstrated that while we can obtain some biologically meaningful results, bulk averaging might lead to significant number of potentially false positive and false negative discoveries.

## Conclusion

I developed a novel computational method called perturb-Met that uses KL divergence to identify metabolite-centric transcriptional changes between two conditions. The main innovation of perturb-Met is considering expression changes in one metabolic enzyme in the context of other enzymes consuming/producing the same metabolite. We demonstrated that perturb-Met can recover literature-supported metabolic changes that are missed by existing methods. We then revealed many neuron subtype-specific metabolic changes in Alzheimer’s disease. The fact that most of these changes enjoy strong literature support suggests that the novel ones uncovered by perturb-Met hold significant promise for future targeted metabolomics experiments and/or enzyme inhibitor experiments. Additionally, the neuron subtype-specific metabolic changes inferred by perturb-Met in human Alzheimer’ disease (**Fig. 3** and **4**) will be a valuable resource for researchers to use when developing targeted metabolomic experiments. We also demonstrated that applying perturb-Met on neuron sub-clusters revealed many literature-supported and novel metabolites that are missed by bulk averaging of all neurons (**Fig. 5**).

I anticipate that perturb-Met will have wide applicability. In the field of immunometabolism, to reveal how different types of immune cells rewire their metabolic networks to fulfill diverse function[60]; in regenerative medicine, to reveal metabolic approaches to promote differentiation and maturation [61,62]; in cancer research, to uncover how metastatic cancer cells adapt their metabolism [ref]. perturb-Met can also be applied to the Tabula Muris Senis project [63], which characterizes age-associated transcriptomic changes at the single cell level across 76 tissue-cell types from 23 tissues[64], providing a glimpse into global scale metabolic changes in aging. It can also be applied to reveal metabolites potentially affected by calorie restriction in aging[65].

I do acknowledge the limitations of perturb-Met. Like Reporter Metabolite analysis [11] and other tools and other methods that use transcriptomic data to infer potential metabolic changes [66,67], it relies on expression data, which is not an ideal proxy for metabolite abundance or fluxes. Thus, it is intended as an exploratory tool to contextualize transcriptomic data and generate more specific hypotheses for further experimental validation.

## Materials and Methods

### Data processing

For bulk RNA-seq, the count table from each study was downloaded, and transcript per million (TPM) values were calculated for each gene. In bulk RNA-seq, only genes expressed above 4 TPM in at least one cell type are considered for further analysis. For single cell RNA-seq, the UMI tables from each study were downloaded. UMI counts in each cell were normalized to the total number of UMIs in each cell. In both types of data, cell type annotations from the original studies were used. The mean expression level for each gene in each cell type is then calculated. In single cell RNA-seq, genes are considered only if (1) their mean expression levels are above 0 in either condition, and (2) they are expressed by at least 10 cells in either condition.

### perturb-Met method

We used the Recon 3D metabolic network. We used the gene-reaction and reaction-metabolite associations annotated in the human metabolic network reconstruction to extract metabolite-gene connections. In this case, all genes encoding enzymes that are involved in the synthesis or degradation of a metabolite are considered together.

#### Detection modules

To detect transcriptional changes, we use two separate modules, each of which will generate a permutation p-value (**Fig. 1**). These two p-values are then combined with Fisher’s method (implemented in the metaSeqR package[15]) to create one p-value for each metabolite.

The first module detects relative compositional changes (Figure 1A). For each metabolite, there are k metabolic genes connected to it (k>=2). The mean expression level of each gene i (i=1, 2, 3,.., k) in condition j (j = 1, 2) is M_i,j_. The sum of all metabolic genes connected to a metabolic gene in condition j is 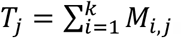. We then normalize m_ij_ by dividing it with T_j_ to obtain 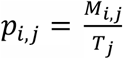. For the first module, the score for each metabolite is then 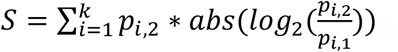.

This scoring scheme will prioritize changes in metabolic genes that are more abundant in condition 2 than other metabolic genes around the same metabolite and exhibit large changes between two conditions.

The first module will miss situations where most genes around a metabolite show similar magnitude of increase/decrease. The second module is intended to capture such situations, and more. In this module, *p*_*i,j*_ and T_j_ are calculated the same way as in module 1. Then, we calculate T_max_ as the maximum of T_1_ and T_2_. We then scale p_ij_ to obtain P_ij_, where 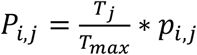. In other words, if the total expression level of all genes in condition 2 is 10-fold higher than in condition 1, P_i,2_ is the same as p_i,2_ because T_2_=T_max_; however, P_i,1_ will be scaled down to 1/10 of p_i,1_. We then assume Pi, j are parameters for a multivariate Dirichlet distribution. We use a formula to calculate KL divergence between these two Dirichlet distributions:

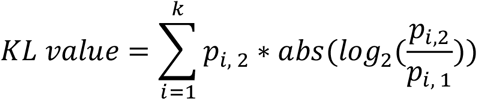

#### Network permutation

Within each module, to assess statistical significance for each metabolite, we randomly sampled k gene expression values from all genes (instead of the actual k metabolic genes connected to the metabolite) and re-calculate scores for each module. p-value is defined as the fraction of times that the scores based on randomly sampled k expression values are equal or larger than the observed scores. The permutation p-value from each method is then combined using Fisher’s method.

#### Cell label permutation

In the AD single cell RNA-seq dataset, we also perform cell label permutation to evaluate whether the observed transcriptional changes around metabolites are due to expression noise. The cell metadata table contains disease condition (AD vs. control) and cluster information (n2, n3, n4, n5) for each cell. We randomly permute the columns of the gene x cell expression matrix to randomize its original correspondence with the cell metadata table. We then run the perturb-Met algorithm to calculate the KL values. We repeat this step 1000 times and only keep metabolites that have cell label permutation p-values of <0.05 (i.e., KL values from cell label permuted expression data exceed the real KL values in less than 50 of 1000 permutations).

## Software Availability

perturb-Met is available both as an R package (https://github.com/yuliangwang/perturb.met).

## Acknowledgements

This project is generously supported by the University of Washington Institute for Stem Cell & Regenerative Medicine.

## Supporting information captions

**Table S1**: List of metabolites identified by perturb-Met as showing significant transcriptional changes of their connected enzymes in APOE4-vs. APOE3-neurons

**Table S2**: List of metabolites identified by perturb-Met as showing significant transcriptional changes of their connected enzymes in APOE4-vs. APOE3-microglia

**Table S3**: List of metabolites identified by perturb-Met as showing significant transcriptional changes of their connected enzymes in neuron sub-cluster n2 between cells from Alzheimer’s Disease and control brain samples.

**Table S4**: List of metabolites identified by perturb-Met as showing significant transcriptional changes of their connected enzymes in neuron sub-cluster n3 between cells from Alzheimer’s Disease and control brain samples.

**Table S5**: List of metabolites identified by perturb-Met as showing significant transcriptional changes of their connected enzymes in neuron sub-cluster n4 between cells from Alzheimer’s Disease and control brain samples.

**Table S6**: List of metabolites identified by perturb-Met as showing significant transcriptional changes of their connected enzymes in neuron sub-cluster n5 between cells from Alzheimer’s Disease and control brain samples.

**Table S7**: List of metabolites identified by perturb-Met as showing significant transcriptional changes of their connected enzymes in the AD-specific n1 neuron sub-cluster and the normal-specific n2 neuron sub-cluster.

**Table S8**: List of metabolites identified by perturb-Met as showing significant transcriptional changes of their connected enzymes in n2-n5 neurons from AD vs. control samples, ignoring sub-cluster identity.

## References

1. GBD 2016 Neurology Collaborators. Global, regional, and national burden of neurological disorders, 1990-2016: a systematic analysis for the Global Burden of Disease Study 2016. Lancet Neurol. 2019;18: 459–480. doi: S1474-4422(18)30499-X [pii].

2. Clarke JR, Ribeiro FC, Frozza RL, De Felice FG, Lourenco MV. Metabolic Dysfunction in Alzheimer’s Disease: From Basic Neurobiology to Clinical Approaches. J Alzheimers Dis. 2018;64: S405–S426. doi: 10.3233/JAD-179911 [doi].

3. Bordbar A, Palsson BO. Using the reconstructed genome-scale human metabolic network to study physiology and pathology. J Intern Med. 2012;271: 131–141. doi: 10.1111/j.1365-2796.2011.02494.x [doi].

4. Lewis NE, Schramm G, Bordbar A, Schellenberger J, Andersen MP, Cheng JK, et al. Large-scale in silico modeling of metabolic interactions between cell types in the human brain. Nat Biotechnol. 2010;28: 1279–1285. doi: 10.1038/nbt.1711 [doi].

5. Stempler S, Yizhak K, Ruppin E. Integrating transcriptomics with metabolic modeling predicts biomarkers and drug targets for Alzheimer’s disease. PloS one. 2014;9: e105383. doi: 10.1371/journal.pone.0105383.

6. Fu H, Hardy J, Duff KE. Selective vulnerability in neurodegenerative diseases. Nat Neurosci. 2018;21: 1350–1358. doi: 10.1038/s41593-018-0221-2 [doi].

7. Zhong S, Zhang S, Fan X, Wu Q, Yan L, Dong J, et al. A single-cell RNA-seq survey of the developmental landscape of the human prefrontal cortex. Nature. 2018;555: 524–528. doi: 10.1038/nature25980 [doi].

8. Ximerakis M, Lipnick SL, Innes BT, Simmons SK, Adiconis X, Dionne D, et al. Single-cell transcriptomic profiling of the aging mouse brain. Nat Neurosci. 2019;22: 1696–1708. doi: 10.1038/s41593-019-0491-3 [doi].

9. Mathys H, Davila-Velderrain J, Peng Z, Gao F, Mohammadi S, Young JZ, et al. Single-cell transcriptomic analysis of Alzheimer’s disease. Nature. 2019;570: 332–337. doi: 10.1038/s41586-019-1195-2.

10. Grubman A, Chew G, Ouyang JF, Sun G, Choo XY, McLean C, et al. A single-cell atlas of entorhinal cortex from individuals with Alzheimer’s disease reveals cell-type-specific gene expression regulation. Nat Neurosci. 2019;22: 2087–2097. doi: 10.1038/s41593-019-0539-4 [doi].

11. Patil KR, Nielsen J. Uncovering transcriptional regulation of metabolism by using metabolic network topology. Proc Natl Acad Sci U S A. 2005;102: 2685.

12. Varemo L, Henriksen TI, Scheele C, Broholm C, Pedersen M, Uhlen M, et al. Type 2 diabetes and obesity induce similar transcriptional reprogramming in human myocytes. Genome Med. 2017;9: 47–2. doi: 10.1186/s13073-017-0432-2 [doi].

13. Wu L, Zhang X, Zhao L. Human ApoE Isoforms Differentially Modulate Brain Glucose and Ketone Body Metabolism: Implications for Alzheimer’s Disease Risk Reduction and Early Intervention. J Neurosci. 2018;38: 6665–6681. doi: 10.1523/JNEUROSCI.2262-17.2018 [doi].

14. Kizuka Y, Kitazume S, Taniguchi N. N-glycan and Alzheimer’s disease. Biochimica et Biophysica Acta (BBA) - General Subjects. 2017;1861: 2447–2454. doi: https://doi.org/10.1016/j.bbagen.2017.04.012.

15. Moulos P, Hatzis P. Systematic integration of RNA-Seq statistical algorithms for accurate detection of differential gene expression patterns. Nucleic Acids Res. 2015;43: e25. doi: 10.1093/nar/gku1273 [doi].

16. Kim J, Basak JM, Holtzman DM. The role of apolipoprotein E in Alzheimer’s disease. Neuron. 2009;63: 287–303. doi: 10.1016/j.neuron.2009.06.026.

17. Lin YT, Seo J, Gao F, Feldman HM, Wen HL, Penney J, et al. APOE4 Causes Widespread Molecular and Cellular Alterations Associated with Alzheimer’s Disease Phenotypes in Human iPSC-Derived Brain Cell Types. Neuron. 2018;98: 1141-1154.e7. doi: S0896-6273(18)30380-5 [pii].

18. Joyce JN, Smutzer G, Whitty CJ, Myers A, Bannon MJ. Differential modification of dopamine transporter and tyrosine hydroxylase mRNAs in midbrain of subjects with parkinson’s, alzheimer’s with parkinsonism, and alzheimer’s disease. Mov Disord. 1997;12: 885–897. doi: 10.1002/mds.870120609.

19. Koch G, Di Lorenzo F, Bonnì S, Giacobbe V, Bozzali M, Caltagirone C, et al. Dopaminergic Modulation of Cortical Plasticity in Alzheimer’s Disease Patients. Neuropsychopharmacology. 2014;39: 2654–2661. doi: 10.1038/npp.2014.119.

20. Cirrito JR, Disabato BM, Restivo JL, Verges DK, Goebel WD, Sathyan A, et al. Serotonin signaling is associated with lower amyloid-beta levels and plaques in transgenic mice and humans. Proc Natl Acad Sci U S A. 2011;108: 14968–14973. doi: 10.1073/pnas.1107411108 [doi].

21. Xie Y, Liu P, Lian Y, Liu H, Kang J. The effect of selective serotonin reuptake inhibitors on cognitive function in patients with Alzheimer’s disease and vascular dementia: focusing on fluoxetine with long follow-up periods. Signal transduction and targeted therapy. 2019;4: 30. doi: 10.1038/s41392-019-0064-7.

22. Riche R, Liao M, Pena IA, Leung KY, Lepage N, Greene NDE, et al. Glycine decarboxylase deficiency-induced motor dysfunction in zebrafish is rescued by counterbalancing glycine synaptic level. JCI Insight. 2018;3: 10.1172/jci.insight.124642. doi: 10.1172/jci.insight.124642 [doi].

23. Ellison DW, Beal MF, Martin JB. Phosphoethanolamine and ethanolamine are decreased in Alzheimer’s disease and Huntington’s disease. Brain Res. 1987;417: 389–392. doi: 0006-8993(87)90471-9 [pii].

24. de Chaves EP, Narayanaswami V. Apolipoprotein E and cholesterol in aging and disease in the brain. Future lipidology. 2008;3: 505–530. doi: 10.2217/17460875.3.5.505.

25. Ambrée O, Richter H, Sachser N, Lewejohann L, Dere E, de Souza Silva, Maria Angelica, et al. Levodopa ameliorates learning and memory deficits in a murine model of Alzheimer’s disease. Neurobiol Aging. 2009;30: 1192–1204. doi: 10.1016/j.neurobiolaging.2007.11.010.

26. Li H, Ye D, Xie W, Hua F, Yang Y, Wu J, et al. Defect of branched-chain amino acid metabolism promotes the development of Alzheimer’s disease by targeting the mTOR signaling. Biosci Rep. 2018;38: BSR20180127. doi: 10.1042/BSR20180127.

27. Chaudhari K, Sumien N, Johnson L, D’Agostino D, Edwards M, Paxton RJ, et al. Vitamin C supplementation, APOE4 genotype and cognitive functioning in a rural-dwelling cohort. J Nutr Health Aging. 2016;20: 841–844. doi: 10.1007/s12603-016-0705-2.

28. Kuiper MA, Visser JJ, Bergmans PL, Scheltens P, Wolters EC. Decreased cerebrospinal fluid nitrate levels in Parkinson’s disease, Alzheimer’s disease and multiple system atrophy patients. J Neurol Sci. 1994;121: 46–49. doi: 10.1016/0022-510x(94)90155-4.

29. Baik SH, Kang S, Lee W, Choi H, Chung S, Kim JI, et al. A Breakdown in Metabolic Reprogramming Causes Microglia Dysfunction in Alzheimer’s Disease. Cell Metab. 2019;30: 493-507.e6. doi: S1550-4131(19)30308-0 [pii].

30. Area-Gomez E, Larrea D, Pera M, Agrawal RR, Guilfoyle DN, Pirhaji L, et al. APOE4 is Associated with Differential Regional Vulnerability to Bioenergetic Deficits in Aged APOE Mice. Scientific Reports. 2020;10: 4277. doi: 10.1038/s41598-020-61142-8.

31. Colton CA, Brown CM, Cook D, Needham LK, Xu Q, Czapiga M, et al. APOE and the regulation of microglial nitric oxide production: a link between genetic risk and oxidative stress. Neurobiology of Aging. 2002;23: 777–785. doi: https://doi.org/10.1016/S0197-4580(02)00016-7.

32. Schlattner U, Tokarska-Schlattner M, Wallimann T. Mitochondrial creatine kinase in human health and disease. Biochimica et Biophysica Acta (BBA) - Molecular Basis of Disease. 2006;1762: 164–180. doi: https://doi.org/10.1016/j.bbadis.2005.09.004.

33. Seminotti B, Zanatta Â, Ribeiro RT, da Rosa MS, Wyse ATS, Leipnitz G, et al. Disruption of Brain Redox Homeostasis, Microglia Activation and Neuronal Damage Induced by Intracerebroventricular Administration of S-Adenosylmethionine to Developing Rats. Mol Neurobiol. 2019;56: 2760–2773. doi: 10.1007/s12035-018-1275-6.

34. Choi YH, Park HY. Anti-inflammatory effects of spermidine in lipopolysaccharide-stimulated BV2 microglial cells. J Biomed Sci. 2012;19: 31. doi: 10.1186/1423-0127-19-31.

35. Lin S, Yin Q, Zhong Q, Lv FL, Zhou Y, Li JQ, et al. Heme activates TLR4-mediated inflammatory injury via MyD88/TRIF signaling pathway in intracerebral hemorrhage. J Neuroinflammation. 2012;9: 46–46. doi: 10.1186/1742-2094-9-46 [doi].

36. Gabbi P, Ribeiro LR, Jessie Martins G, Cardoso AS, Haupental F, Rodrigues FS, et al. Methylmalonate Induces Inflammatory and Apoptotic Potential: A Link to Glial Activation and Neurological Dysfunction. J Neuropathol Exp Neurol. 2017;76: 160–178. doi: 10.1093/jnen/nlw121 [doi].

37. Mills EL, Kelly B, Logan A, Costa ASH, Varma M, Bryant CE, et al. Succinate Dehydrogenase Supports Metabolic Repurposing of Mitochondria to Drive Inflammatory Macrophages. Cell. 2016;167: 457-470.e13. doi: S0092-8674(16)31162-X [pii].

38. Kikusui T, Shimozawa A, Kitagawa A, Nagasawa M, Mogi K, Yagi S, et al. N-Acetylmannosamine Improves Object Recognition and Hippocampal Cell Proliferation in Middle-Aged Mice. Biosci Biotechnol Biochem. 2012;76: 2249–2254. doi: 10.1271/bbb.120536.

39. Tokizane K, Konishi H, Makide K, Kawana H, Nakamuta S, Kaibuchi K, et al. Phospholipid localization implies microglial morphology and function via Cdc42 in vitro. Glia. 2017;65: 740–755. doi: 10.1002/glia.23123 [doi].

40. Xu J, Drew PD. 9-Cis-retinoic acid suppresses inflammatory responses of microglia and astrocytes. J Neuroimmunol. 2006;171: 135–144. doi: S0165-5728(05)00427-3 [pii].

41. Stojakovic A, Trushin S, Sheu A, Khalili L, Chang S, Li X, et al. Partial inhibition of mitochondrial complex I attenuates neurodegeneration and restores energy homeostasis and synaptic function in a symptomatic Alzheimer’s mouse model. bioRxiv. 2020: 2020.07.01.182428. doi: 10.1101/2020.07.01.182428.

42. Zhang L, Zhang S, Maezawa I, Trushin S, Minhas P, Pinto M, et al. Modulation of mitochondrial complex I activity averts cognitive decline in multiple animal models of familial Alzheimer’s Disease. EBioMedicine. 2015;2: 294–305. doi: 10.1016/j.ebiom.2015.03.009 [doi].

43. Shi A, Petrache AL, Shi J, Ali AB. Preserved Calretinin Interneurons in an App Model of Alzheimer’s Disease Disrupt Hippocampal Inhibition via Upregulated P2Y1 Purinoreceptors. Cereb Cortex. 2020;30: 1272–1290. doi: 10.1093/cercor/bhz165.

44. Kracun I, Kalanj S, Talan-Hranilovic J, Cosovic C. Cortical distribution of gangliosides in Alzheimer’s disease. Neurochem Int. 1992;20: 433–438. doi: 10.1016/0197-0186(92)90058-y [doi].

45. Amaro M, Sachl R, Aydogan G, Mikhalyov II, Vacha R, Hof M. GM1 Ganglioside Inhibits beta-Amyloid Oligomerization Induced by Sphingomyelin. Angew Chem Int Ed Engl. 2016;55: 9411–9415. doi: 10.1002/anie.201603178 [doi].

46. Shin M, Choi M, Chae H, Kim J, Kim H, Kim K. Ganglioside GQ1b ameliorates cognitive impairments in an Alzheimer’s disease mouse model, and causes reduction of amyloid precursor protein. Scientific Reports. 2019;9: 8512. doi: 10.1038/s41598-019-44739-6.

47. Chung H, Park K, Jang HJ, Kohl MM, Kwag J. Dissociation of somatostatin and parvalbumin interneurons circuit dysfunctions underlying hippocampal theta and gamma oscillations impaired by amyloid beta oligomers in vivo. Brain Struct Funct. 2020;225: 935–954. doi: 10.1007/s00429-020-02044-3 [doi].

48. Verret L, Mann EO, Hang GB, Barth AM, Cobos I, Ho K, et al. Inhibitory interneuron deficit links altered network activity and cognitive dysfunction in Alzheimer model. Cell. 2012;149: 708–721. doi: 10.1016/j.cell.2012.02.046 [doi].

49. Cui X, Zuo P, Zhang Q, Li X, Hu Y, Long J, et al. Chronic systemic D-galactose exposure induces memory loss, neurodegeneration, and oxidative damage in mice: protective effects of R-alpha-lipoic acid. J Neurosci Res. 2006;83: 1584–1590. doi: 10.1002/jnr.20845 [doi].

50. Stopschinski BE, Holmes BB, Miller GM, Manon VA, Vaquer-Alicea J, Prueitt WL, et al. Specific glycosaminoglycan chain length and sulfation patterns are required for cell uptake of tau versus α-synuclein and β-amyloid aggregates. J Biol Chem. 2018;293: 10826–10840. doi: 10.1074/jbc.RA117.000378 [doi].

51. Choi JW, Shin CY, Choi MS, Yoon SY, Ryu JH, Lee JC, et al. Uridine protects cortical neurons from glucose deprivation-induced death: possible role of uridine phosphorylase. J Neurotrauma. 2008;25: 695–707. doi: 10.1089/neu.2007.0409 [doi].

52. Xu Y, Zhao M, Han Y, Zhang H. GABAergic Inhibitory Interneuron Deficits in Alzheimer’s Disease: Implications for Treatment. Front Neurosci. 2020;14: 660. doi: 10.3389/fnins.2020.00660 [doi].

53. Mobin MB, Gerstberger S, Teupser D, Campana B, Charisse K, Heim MH, et al. The RNA-binding protein vigilin regulates VLDL secretion through modulation of Apob mRNA translation. Nat Commun. 2016;7: 12848. doi: 10.1038/ncomms12848 [doi].

54. Chiu DS, Oram JF, LeBoeuf RC, Alpers CE, O’Brien KD. High-density lipoprotein-binding protein (HBP)/vigilin is expressed in human atherosclerotic lesions and colocalizes with apolipoprotein E. Arterioscler Thromb Vasc Biol. 1997;17: 2350–2358. doi: 10.1161/01.atv.17.11.2350 [doi].

55. Yang C, Wang X, Wang J, Wang X, Chen W, Lu N, et al. Rewiring Neuronal Glycerolipid Metabolism Determines the Extent of Axon Regeneration. Neuron. 2020;105: 276-292.e5. doi: S0896-6273(19)30882-7 [pii].

56. Schedin-Weiss S, Winblad B, Tjernberg LO. The role of protein glycosylation in Alzheimer disease. FEBS J. 2014;281: 46–62. doi: 10.1111/febs.12590 [doi].

57. DeWitt DA, Silver J, Canning DR, Perry G. Chondroitin sulfate proteoglycans are associated with the lesions of Alzheimer’s disease. Exp Neurol. 1993;121: 149–152. doi: S0014-4886(83)71081-2 [pii].

58. Yang S, Hilton S, Alves JN, Saksida LM, Bussey T, Matthews RT, et al. Antibody recognizing 4-sulfated chondroitin sulfate proteoglycans restores memory in tauopathy-induced neurodegeneration. Neurobiol Aging. 2017;59: 197–209. doi: S0197-4580(17)30257-9 [pii].

59. Srinivasan K, Friedman BA, Larson JL, Lauffer BE, Goldstein LD, Appling LL, et al. Untangling the brain’s neuroinflammatory and neurodegenerative transcriptional responses. Nat Commun. 2016;7: 11295. doi: 10.1038/ncomms11295 [doi].

60. Ganeshan K, Chawla A. Metabolic regulation of immune responses. Annu Rev Immunol. 2014;32: 609–634. doi: 10.1146/annurev-immunol-032713-120236 [doi].

61. Sperber H, Mathieu J, Wang Y, Ferreccio A, Hesson J, Xu Z, et al. The metabolome regulates the epigenetic landscape during naive-to-primed human embryonic stem cell transition. Nat Cell Biol. 2015;17: 1523–1535. doi: 10.1038/ncb3264.

62. Ellen Kreipke R, Wang Y, Miklas JW, Mathieu J, Ruohola-Baker H. Metabolic remodeling in early development and cardiomyocyte maturation. Semin Cell Dev Biol. 2016;52: 84–92. doi: 10.1016/j.semcdb.2016.02.004 [doi].

63. Tabula Muris Consortium. A single-cell transcriptomic atlas characterizes ageing tissues in the mouse. Nature. 2020;583: 590–595. doi: 10.1038/s41586-020-2496-1 [doi].

64. , Pisco AO, McGeever A, Schaum N, Karkanias J, Neff NF, et al. A Single Cell Transcriptomic Atlas Characterizes Aging Tissues in the Mouse. bioRxiv. 2020: 661728. doi: 10.1101/661728.

65. Ma S, Sun S, Geng L, Song M, Wang W, Ye Y, et al. Caloric Restriction Reprograms the Single-Cell Transcriptional Landscape of Rattus Norvegicus Aging. Cell. 2020;180: 984-1001.e22. doi: 10.1016/j.cell.2020.02.008.

66. Schramm G, Wiesberg S, Diessl N, Kranz AL, Sagulenko V, Oswald M, et al. PathWave: discovering patterns of differentially regulated enzymes in metabolic pathways. Bioinformatics. 2010;26: 1225–1231. doi: 10.1093/bioinformatics/btq113 [doi].

67. Blazier AS, Papin JA. Integration of expression data in genome-scale metabolic network reconstructions. Front Physiol. 2012;3: 299.

